# ZeBox: A novel non-intrusive continuous-use technology to trap and kill airborne microbes

**DOI:** 10.1101/2021.08.02.454789

**Authors:** Kruttika S. Phadke, Deepak G. Madival, Janani Venkataraman, Debosmita Kundu, K. S. Ramanujan, Nisha Holla, Jaywant Arakeri, Gaurav Tomar, Santanu Datta, Arindam Ghatak

## Abstract

Preventing nosocomial infection is a major unmet need of our times. Existing air decontamination technologies suffer from demerits such as toxicity of exposure, species specificity, noxious gas emission, environment-dependent performance and high power consumption. Here, we present a novel technology called “ZeBox” that transcends the conventional limitations and achieves high microbicidal efficiency. In ZeBox, a non-ionizing electric field extracts naturally charged microbes from flowing air and deposits them on engineered microbicidal surfaces. The surface’s three dimensional topography traps the microbes long enough for them to be inactivated. The electric field and chemical surfaces synergistically achieve rapid inactivation of a broad spectrum of microbes. ZeBox achieved near complete kill of airborne microbes in challenge tests (5-9 log reduction) and >90% efficiency in a fully functional stem cell research facility in the presence of humans. Thus, ZeBox fulfills the dire need for a real-time, continuous, safe, trap-and-kill air decontamination technology.

## 1 Introduction

Microbial load (bacteria, viruses, spores and fungi) in our living, working and hospital space must be reduced to mitigate the transmission of airborne infections. As per CDC (Center for Disease Control, USA)’s recommendation (https://www.cdc.gov/niosh/topics/hierarchy/default.html), eliminating microbes at the source as and when produced is the first line of defense against spread of infections. Filtration, electrostatic precipitation, bactericidal gas spraying, ultra-violet germicidal irradiation (UVGI, employing 254 ∼ nm radiation), plasma discharge and photo-catalytic oxidation (PCO) are the currently available air decontamination technologies [1]. While some are microbicidal, others only trap microbes. Filtration [2] and electrostatic precipitation [3] belong to the latter category. Microbes trapped inside filters can multiply in situ [4, 5, 6, 7, 8]; such filters are detrimental to indoor air quality and hazardous during their disposal. They also offer high flow resistance which translates to high operating power consumption [9, 10]. Electrostatic precipitation uses electric field to attract and trap aerosols pre-charged by corona discharge, but which produces noxious gases like ozone [3, 11]. Its microbicidal action is dubious; in fact electrostatic bioaerosol samplers capture microbes that remain viable [12, 13, 14]. However, because of its low flow resistance, it consumes less power per unit of clean air delivered compared to filtration [3]. Filters made of anti-bacterial fibers have also been developed [15, 16, 17, 18, 19, 20] but their performance remains be proven under realistic indoor conditions.

Bactericidal gas spraying, UVGI, plasma discharge and PCO are microbicidal technologies. Although bactericidal gases and UVGI can sterilize an entire room, they cannot be deployed in human presence. UVGI is used to sterilize upper room air and air circulating through ventilation ducts. However, microbicidal action of UVGI depends on environmental parameters such as humidity [21, 22, 23], is species-specific [24] and requires a minimum duration of exposure to microbes [25]. Exposure of humans to UVGI (due to faulty design, deployment or use of UVGI devices) can damage their eyes and skin [26, 27, 28, 29]. UVGI is used to kill microbes trapped on a filter’s surface [30, 31] but then it cannot reach microbes residing beneath the surface. Plasma discharge [32] and PCO [34, 35] both generate ions and/or reactive species, respectively using gas discharge and reaction with an irradiated catalyst. However, they also generate NO_*X*_ and ozone [1] and additional methods are necessary to mitigate them [33]. In PCO, convection of gas to the catalyst and the subsequent adsorption, reaction and release of reactive species into the bulk flow is the bottleneck process [36], which results in low clean air delivery rates [1].

Given the importance of eliminating airborne infection, a technology that is safe, suitable for continuous use and efficient against a wide variety of airborne microbes is desirable. Here, we describe such a novel technology called “ZeBox”; the name derives from the **Ze**ta-potential possessed by microbes, which property is pivotal in trapping them inside the **Box**-shaped device. In the following, we discuss the working mechanism of ZeBox and demonstrate its efficacy in chamber tests and field studies against a variety of microbes.

## 2 Results

### Electrode plates with engineered chemical surfaces form the kill cassette

A row of flat plate electrodes (10.9 cm × 30 cm) with alternating polarity are assembled inside a cuboid shaped box with open ends for transmitting flow. A three dimensional hydrocellular microbicidal composite material (US patent no. US 9566363B2, licensed) is layered on to the electrodes. A non-ionizing 3 kV/cm electric field is set up between electrodes by applying direct-current voltage between them. Microbes are trapped and killed inside this “kill cassette”. Axial fans pull microbe laden ambient air through the kill cassette and between the electrode-plates, as shown schematically in figure 1.

**Figure 1:**
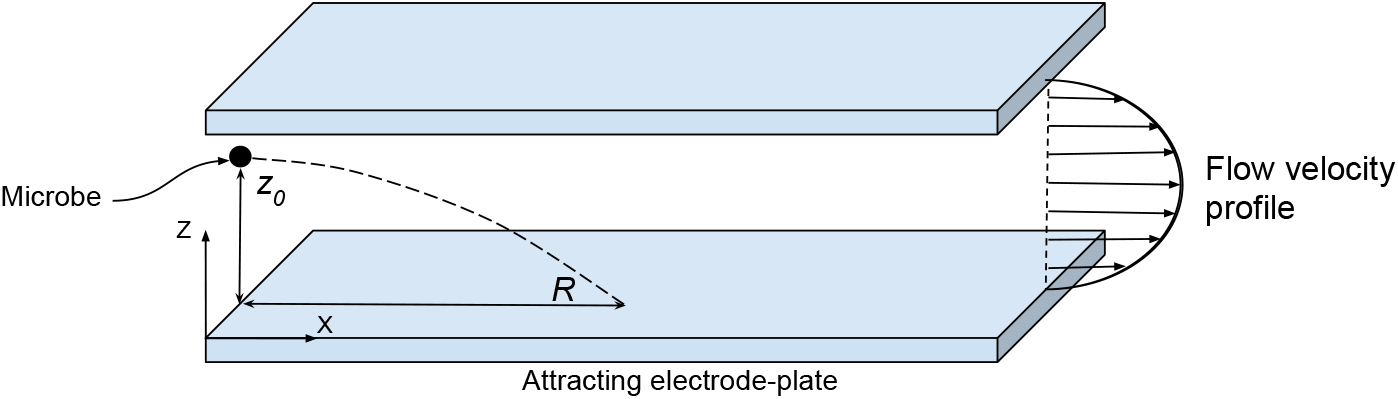
Microbe in a flow subject to transverse electric field. A charged microbe deviates from the flow direction due to the electric field between electrode-plates.

### Electric field extracts charged microbes from the flow

Microbes are naturally charged [38, 39]; therefore, in an electric field, they are impelled towards the electrode of opposite polarity. Figure 1 depicts this process schematically. Here, *X*-axis points along the flow and *Z*-axis points away from the attracting electrode. A microbe initially at distance *z*_0_ from the attracting electrode travels a distance *R* in the streamwise direction, called its “range”, as it descends to *z* = 0. Whether or not the microbe hits the electrode depends on its length, the microbe’s initial distance *z*_0_, strength of the electric field, charge on the microbe and the type of flow (laminar or turbulent). The Reynolds number for the flow between electrodes in ZeBox is ∼ 10^3^ and a rectangular duct flow (or even plane Poiseuille flow) undergoes transition at this Reynolds number and could be turbulent [40, 41]. Analyzing microbe’s motion in a turbulent flow is difficult because of its complicated, stochastic nature; supplementary information S1 analyzes microbe’s motion and its maximum range in a laminar flow instead.

Earlier studies on resuspension of dust from flat surfaces due to a flow show that, whenever the hydrodynamic force and torque exerted by the flow exceed those that keep the particles attached to the surface (for example, Van der Waals force), the particles can either detach by lifting off or slide and roll on the surface [42, 43]. In our case, lifting off of microbes from the electrode is unlikely due to the strong electric field, but they can nevertheless slide and roll and thus escape away due to the electrode’s finite length (refer figure 2). Since the microbicidal surface requires a minimum duration of contact to inactivate microbes depending on how sensitive or hardy it is, a fraction of the deposited microbes could escape while still viable. Therefore, the ability of the microbicidal surface to trap and hold microbes until they are inactivated becomes important.

**Figure 2:**
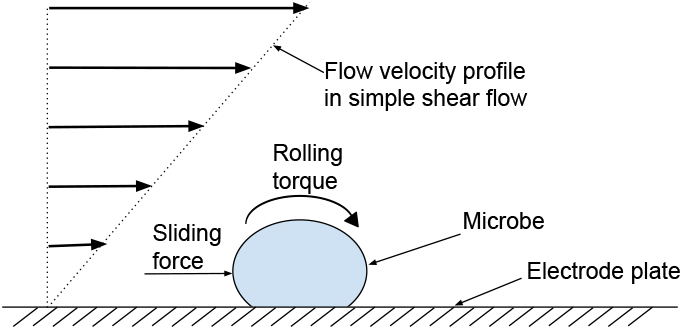
Microbe sliding and rolling on a flat surface. Microbes can slide and roll over a flat surface due to hydrodynamic force and torque exerted by the flow.

### Three dimensional topography of the microbicidal surface traps the microbe

The microbicidal surface employed in ZeBox has a highly uneven topography at the microbial scale, populated with well-like depressions to trap and hold microbes. Figures 3a and 3b show the scanning electron microscope (SEM) images of the surface at different magnifications appropriate to the microbial scale. Figure 3c shows streamlines in a numerically simulated two dimensional flow (using OpenFOAM-7) over a surface with square shaped wells, to qualitatively illustrate the kind of flow obtained over an uneven topography. A simple shear flow was imposed on the flow domain (refer figure 3c) by moving its uppermost boundary horizontally at constant speed. The flow Reynolds number based on the imposed shear rate and the dimension of the square-shaped well is ∼ 10^-5^, which is appropriate to the flow in the neighborhood of the microbicidal surface in ZeBox. The important feature of the flow for our purpose is the recirculating region set up within the wells, in which the streamlines of the flow form closed loops. This feature is quite general for a flow over an uneven topography and which presumably enhances the efficacy of the microbicidal surface further in regard to trapping microbes. Once the microbe falls into one of the wells, brought there either in the course of its rolling over the surface or directly by the electric field, the recirculating flow can confine it to the well for a sufficiently long duration.

**Figure 3:**
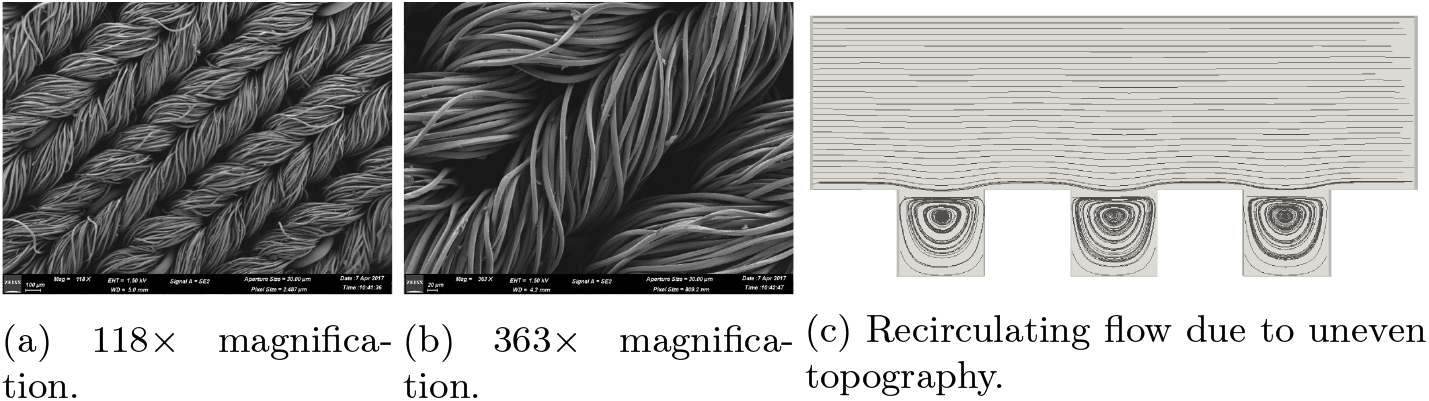
Topography of the microbicidal surface at microbial scale. SEM photographs revealing the highly uneven topography of microbicidal surface and the expected flow patterns over it.

Table 1 shows the efficacy of microbicidal surfaces (in terms of log_10_ reduction, where *n*-log_10_ reduction implies reduction in the initial microbial load by a factor of 10^*n*^) with different topographies, which we call 2-D and 3-D surfaces, in flow experiments. A 2-D surface is a single layer of cotton fabric while a 3-D surface is a multilayered 90:10 polyethylene : cotton fabric. In the presence of electric field, 3-D microbicidal surface performs better than the 2-D surface. When the electric field is absent, the microbes are not extracted from the flow and hence both surfaces perform similarly.

**Table 1:**
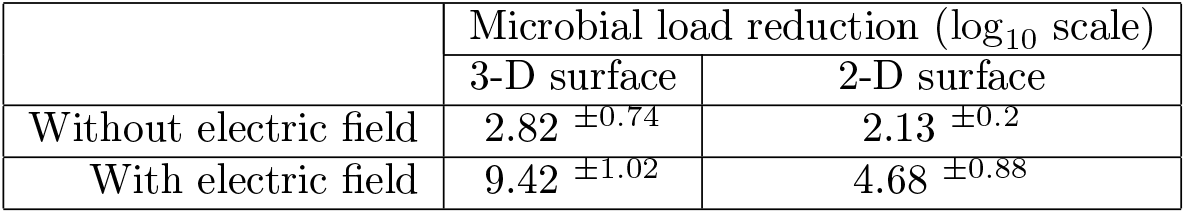
Effect of surface topography on microbicidal action. Log_10_-reduction in viable microbial load (*E. coli*) achieved by ZeBox with 3-D and 2-D microbicidal surfaces in 10 minutes. Applied electric field = 3 kV/cm. Superscripts show standard deviation.

| | Microbial load reduction ($\log_{10}$ scale) | |
| --- | --- | --- |
|  | 3-D surface | 2-D surface |
| Without electric field | $2.82 \pm 0.74$ | $2.13 \pm 0.2$ |
| With electric field | $9.42 \pm 1.02$ | $4.68 \pm 0.88$ |

### Electric field and chemical microbicidal-surfaces synergistically achieve rapid inactivation of microbes

In contrast to electrostatic precipitators, the applied electric field in ZeBox plays two roles: it pulls microbes from the flow on to the microbicidal surface and then accelerates their subsequent inactivation. Table 2 shows log_10_-reduction in the microbial load in spot experiments, with 3 kV/cm electric field applied between electrodes. The microbicidal surface achieves the highest reduction in microbial load in the presence of the electric field. Quaternary ammonium compounds (QAC) are membrane-active agents which inactivate microbes by targeting their cytoplasmic membrane [46, 47, 48, 49], but first, they must breach the outer cell wall. In the present design, QAC is tethered to the 3-D surface by long flexible chains, which presumably helps the QAC to orient itself to puncture holes in the microbe. The external electric field increases the trans-membrane voltage of the cell above its resting value, leading to an electric current that presumably flows through these pores as they form the path of least resistance. This current flow may be analogous to the electroporation of bacteria in which the pores formed in the cell wall are stabilized [50]. The intracellular components then leak from the pores, as is seen in the SEM pictures. This process leads to the irreversible killing of the cells. Therefore, the chemical surface in tandem with the electric field displays an enhanced electro-chemical microbicidal action compared to what they would have achieved separately.

**Table 2:** Effect of applied electric field on the efficacy of microbici-dal surface. Effect of 3 kV/cm electric field on the log_10_-reduction in viable microbial load (*E. coli*) over the microbicidal surface in spot experiments. Su-perscripts show standard deviation.

| Time (mins) | Microbial load reduction ($\log_{10}$ scale) | |
| --- | --- | --- |
|  | With electric field | Without electric field |
| 2 | $3.00 \pm 0.39$ | $0.87 \pm 0.44$ |
| 5 | $5.71 \pm 0.19$ | $1.86 \pm 0.78$ |
| 10 | $8.83 \pm 0.69$ | $2.56 \pm 1.17$ |

### ZeBox rapidly reduces microbial load in chamber tests

The capability of ZeBox to decontaminate a closed space containing airborne microbes was determined by challenge tests [51]. A broad spectrum of microorganisms was employed in the test – standard gram-positive and gram-negative bacteria of ESKAPE group (*Escherichia coli, Staphylococcus aureus, Pseudomonas aeruginosa*), mycobacterium species (*Mycobacterium smegmatis*), fungal species (*Aspergillus fumigatus* spores and *Candida albicans*) and virus (PhiX 174 coliphage and MS2 coliphage). Among these, MS2 virus is an accepted surrogate for the SARS-CoV2 virus [52, 53]. Figure 4 shows the collated data on the variation in log_10_ microbial load (*n*-log_10_ microbial load equals 10^*n*^ microbes) over time after ZeBox was turned on. ZeBox proves to be extremely effective in rapidly decreasing the viable microbial load in a closed space. It achieved 9.9 log_10_-reduction (i.e. 99.999999999% reduction) of *E. coli* in 10 minutes (*n* log_10_-reduction equals reduction by a factor of 10^*n*^). For other microbes ZeBox brought about 5 to 9 log_10_-reduction (i.e. 99.999-99.9999999% reduction) of the viable microbial load.

**Figure 4:**
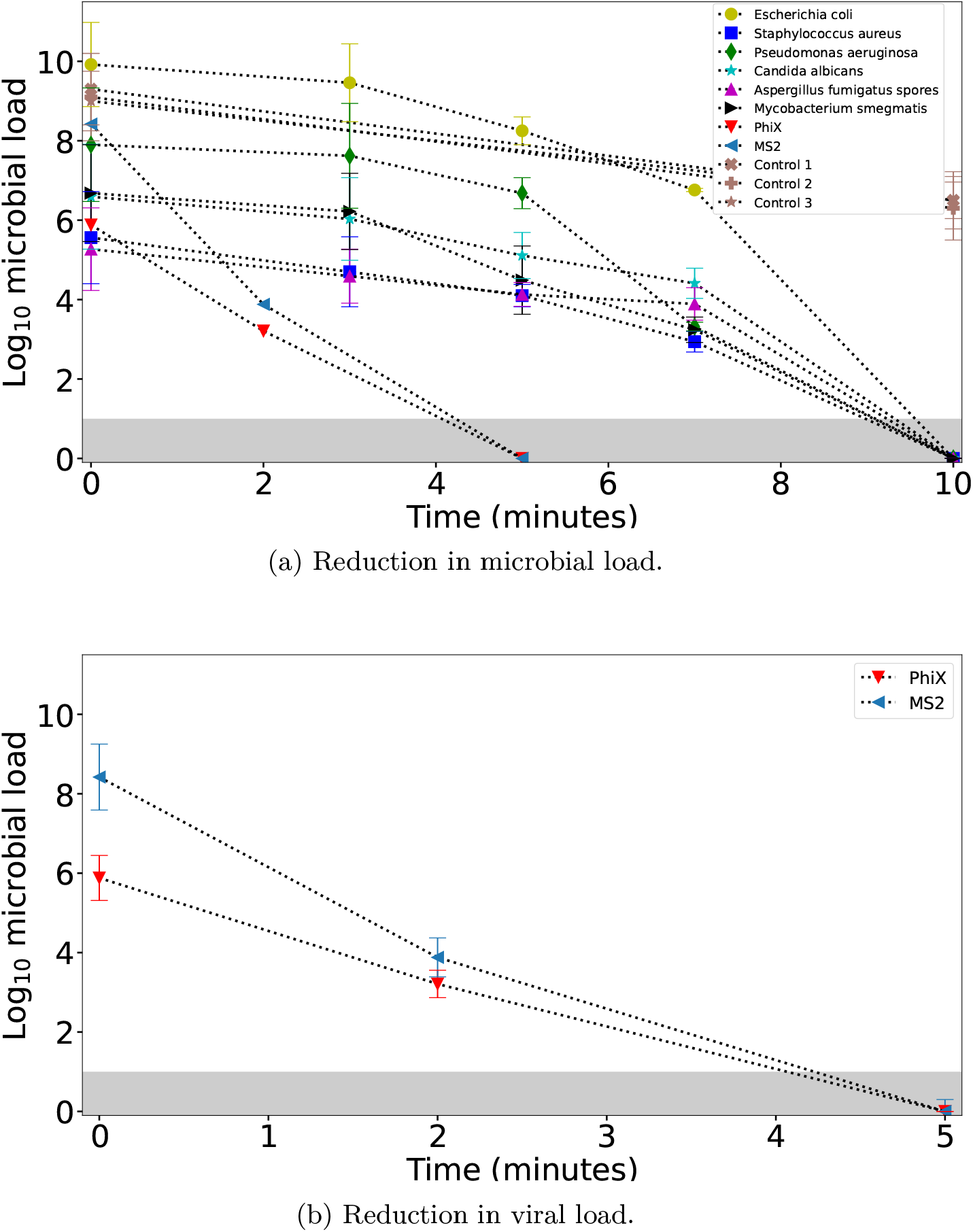
Reduction of microbial load in test chamber. Variation of the log_10_ microbial load over time after ZeBox is turned on in the test chamber. The shaded region indicates limit of detection (LoD). Control 1, 2, 3 refer to control experiments employing respectively microbicidal surface without electric field, control surface with electric field and control surface without electric field.

### SEM images of microbicidal action

Scanning electron microscopy (SEM) studies were done to see how microbes trapped on the microbicidal surface are killed. *E. coli* and *A. fumigatus* spores were chosen because they form two extremes on the scale of sensitivity, with spores being hardy. Figure 5a and 5e show the microbes in control conditions. Due to electro-chemical action at the three dimensional microbicidal surface, their cell membrane undergoes morphological changes followed by complete degradation. Figure 5b and 5c, obtained after 5 minutes of contact, reveals puncturing and blebbing of the *E. coli* cell membrane. Ultimately, the cells burst and their intracellular contents spill out (figure 5d and 5f) signaling a complete degradation of the microbes.

**Figure 5:**
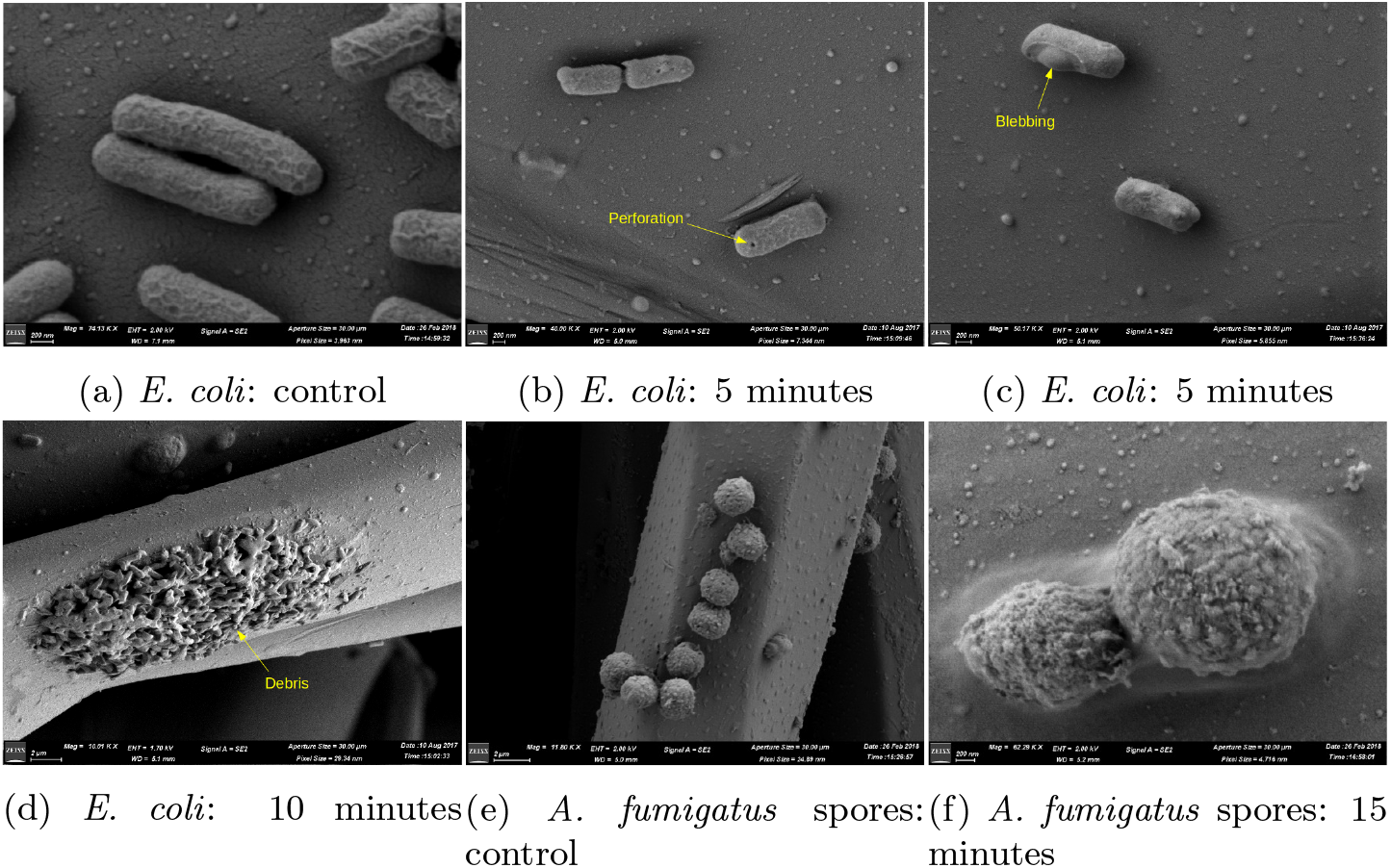
Microbicidal action of the chemical surface. SEM images show-ing microbes on the microbicidal surface being killed by electroporation.

### ZeBox reduces microbial load in open room

ZeBox’s performance was also tested in a real life setting, i.e. in a room with constant influx of microbes from outside or due to internal sources. A working tissue culture laboratory in a building with central air-conditioning, but without High Efficiency Particulate Air (HEPA) filters, was chosen for the purpose. Figure 6a shows the schematic plan-view of the lab and the measurement locations. The working people in the lab were the primary source of microbial contamination. Figure 6b shows that the microbial load at location-03 where tissue culture work was carried out was >3000 CFU/m^3^ initially. ZeBox reduced the microbial load in the lab to ∼ 10 CFU/m^3^ within about 3 hours after it was turned on. This low level was consistently maintained so long as ZeBox was operational. When it was turned off at day 10, the microbial load rebounded to its original level. During its operation, ZeBox effectively decontaminated a zone of dimensions ∼ 10 feet × 10 feet (refer figure 6a), which demonstrates its potential to decontaminate a smaller region of interest in a relatively large open room, with continuous movement of personnel and without needing physical partitions.

**Figure 6:**
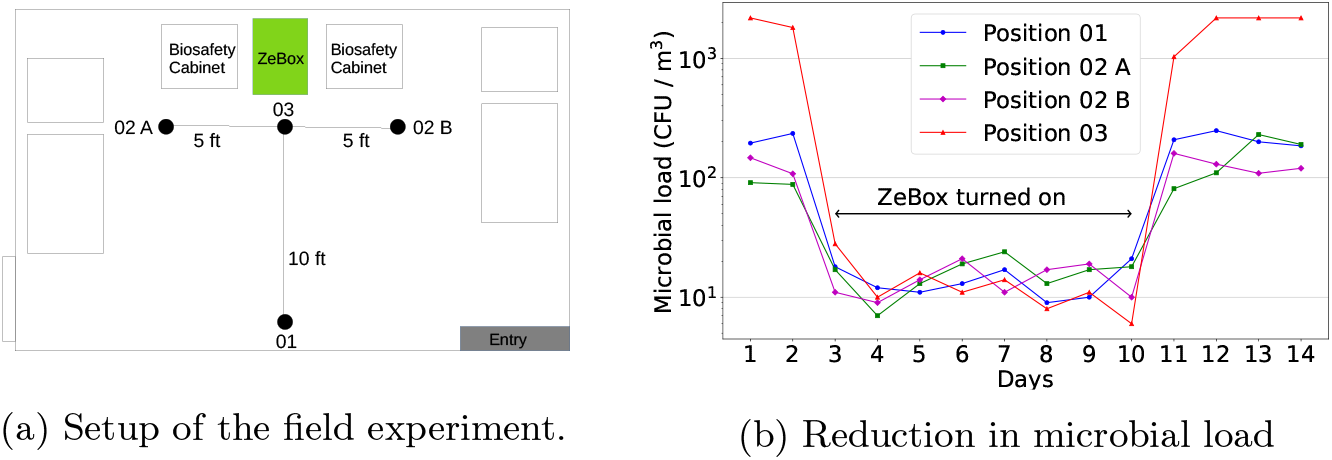
Field performance of ZeBox. ZeBox reduces the microbial load in an open room. Measurement locations are indicated by filled circles in the schematic.

### ZeBox does not produce ozone

Since ZeBox employs non-ionizing electric field, it does not produce ozone (verified in standardized laboratory tests, data not shown here). This is an immense advantage over conventional microbicidal technologies such as plasma and PCO. Also, it consumes <20 Watt-hour of electric energy during its operation.

## Discussion

ZeBox technology exploits the fact that microbes (bacteria, viruses, spores and fungi) are naturally charged and therefore can be readily manipulated by an electric field. Using a non-ionizing electric field, microbicidal surfaces with three dimensional topography and electro-chemical kill mechanism, ZeBox achieves significantly higher microbicidal rate compared to other technologies.

Knowing the total reduction in microbial load, as shown in figure 4, is inadequate to gauge ZeBox’s efficacy because any level of decontamination may be achieved given sufficient time. Therefore, an overall microbicidal efficiency must be determined while factoring in the time of operation as well as the volume of the room being decontaminated. Towards this end, we may think in terms of the number of nominal air changes in a room achieved in a given duration and the consequent reduction in microbial load for each air change. In time *t, Qt/V* number of nominal air changes is achieved, where *Q* is the air flow rate through ZeBox and *V* is the volume of the room. If *η* is the corresponding microbicidal efficiency, then *N*_0_ initial number of viable microbes in the room decreases to *N* = *N*_0_(1-*η*)^*Qt/V*^ after time *t*. Using this formula and the latest-time data from figure 4 whose ordinate is log_10_ *N*, we may back-calculate *η* for a specified time duration. The microbicidal efficiency of ZeBox lies in the range of 83-99 % for all the tests. Considering the variety of sensitive and hardy microbes employed, ZeBox is about equally effective against all of them. Supplementary information S2 provides a theoretical estimation of the microbicidal efficiency of ZeBox and shows that the efficiency deduced from experimental data is aligned with it.

Airborne microbes of size *<* 2 *µ*m can remain suspended in air for several hours before settling down and therefore must be inactivated to reduce the transmission of infections. ZeBox technology presents a universal solution because:

- Freely floating microbes are trapped and killed with high efficiency, eliminating the possibility of future growth.
- The airflow is parallel to antimicrobial surfaces with almost no resistance; therefore, unlike HEPA filters, it has low energy utilization.
- There are no chemical emissions or production of free radicals or ozone; the technology is safe for continuous use in the presence of humans and animals.
- It is equally effective for different varieties of sensitive and hardy microbes.

## Materials and methods

### Challenge tests

#### A. Test setup

An air-sealed test chamber of dimensions 3 ft × 4 ft × 3 ft (approximately 1000 liters in volume) was built with multiple sampling and nebulization ports. The environmental parameters such as relative humidity and temperature could be monitored using a probe located inside the chamber. During experiments, various microorganisms were aerosolized using a 6-jet collision nebulizer (MESA LABS, BGI) into the chamber, and the device efficiency was monitored by collecting and measuring microbial concentration at different time intervals. A second test chamber of dimensions 3ft x 2.5 ft x 1 ft (approximately 220 liters in volume) placed inside a biosafety cabinet, with similar aerosolization and sampling port configuration, was used for tests with viruses.

#### B. Cultivation of test microorganisms

To validate the efficiency of the decontamination device, *Escherichia coli* (MTCC 40), *Pseudomonas aeruginosa* (MTCC 424), *Staphylococcus aureus* (MTCC 96), *Candida albicans* (MTCC 584), *Aspergillus fumigatus* (MTCC 2544), *Mycobacterium smegmatis* (MTCC 6), MS2 coliphage (ATCC 15597-B1) and PhiX 174 coliphage (ATCC 13706-B1) were used. For growing *Escherichia coli, Pseudomonas aeruginosa* and *Staphylococcus aureus*, Luria broth was used. For growing *Candida albicans*, Potato dextrose broth was used, while for *M. smegmatis*, Middlebrook 7H9 broth was used. For enumeration of *E*.*coli*, samples were plated on Luria Bertani agar; Cetrimide agar was used as a selective for the growth and isolation of *Pseudomonas aeruginosa*. Cetrimide inhibits the growth of many microorganisms while allowing *Pseudomonas aeruginosa* to develop typical colonies. For quantification of *Staphylococcus*, Mannitol Salt Agar plates were used. *Candida albicans* and *Aspergillus fumigatus* spores were enumerated using Rose-Bengal Chloramphenicol Agar plates. Coliphages were cultivated using standard method described in ATCC manual. For all microbiological nutrient media were manufactured by HiMedia Laboratories, India unless mentioned otherwise.

#### C. Aerosolization of test microbes

A 6-jet Collison nebulizer (MESA LABS, BGI) was used to aerosolize the test microbes into the test chamber. Dry air from a compressed air cylinder at a pressure of 10 psi was used to operate the nebulizer. The nebulizer produces bioaerosols of a 2-5 *µ*m diameter that allows them to float in the air present in the test chamber for a definite period. The length of the nebulization period varied depending on the type of experiment and microorganism, but typically ranged between 30-40 minutes.

#### D. Sampling of air for viable microbes

The airborne survival of the test microbe and the activity of the air decontamination devices were determined by collecting the air from the chamber at the rate of 12.5 liter/min using SKC biosampler [54], filled with sterile buffer (1x Phosphate buffer saline, pH 7.2). Collected samples were analyzed to understand the quantity of viable microorganism present by diluting and plating them onto suitable growth media. The plated samples were incubated at 37±2 ^0^C for bacteria and 25±2 ^0^C for fungal species and allowed to grow for 18-48 hours as mentioned in the ATCC/MTCC manual, individual colonies were enumerated, and the final concentration of the microbial load was calculated thereafter. For enumerating coliphages collected from the chamber, Double agar overlay method was used for subsequent plaque assay [55]. *E. coli* ATCC 15597 and *E. coli* ATCC 13706 were used as a host in plaque assays for MS2 and PhiX174, respectively. Plaques were counted after 24 hour incubation at 37±2 ^0^C.

#### E. Spot experiments

*E. coli* cells were grown in the standard medium. A known titre of cells were spotted onto a 25 mm^2^ surface and incubated for various time duration, both with and without electric field. Surfaces were resuspended in 500 *µ*l of sterile 1X PBS, which was then plated on standard agar plates to enumerate the microbes.

#### F. Limit of Detection

Microbial enumeration is guided by two parameters, Limit of Detection (LOD) and Limit of Quantification (LOQ). For the present assays used to quantify the microbial load inside the test chamber, the LOD was 10 CFU for bacterial and fungal load and 5 PFU for viral load. However, LOD is always less than LOQ [56]. In many of our experimental analysis, post operating ZeBox device, the microbial detected numbers were in around LOD and hence, the exact LOQ was often indeterminant.

#### G. SEM analysis of trapped microbes to decipher the mechanism of kill

3D surfaces were stripped off from the electrode plates post operating the device against E.coli under challenge test under various time course, and treated with 2.5% glutaraldehyde in 0.1 M phosphate buffer (pH 7.2) for 24 hrs at 4 ^0^C. The samples were dehydrated in series of graded ethanol solutions and subjected to critical point drying with CPD unit. The analyzed samples were mounted over the stud with double-sided carbon conductivity tape, and a thin layer of gold coat over the samples was done by using an automated sputter coater (EMITECK K550X Sputter Coater from EM Scientific Services) for 3 minutes and analyzed under Field Emission Scanning Electron Microscope (MERLIN Compact VP from M/s.Carl Ziess). The set parameters were: Working Distance= 5-6 mm, EHT range= 2-4 kV, Range of Magnification= 70 KX, detectors=SE2 And InLens, machine under high vacuum.

### Field tests

#### H. Air sample collection

A working tissue culture laboratory in a national stem cell research facility was chosen for study. This laboratory was situated in a building which had central airconditioning but the absence of a HEPA-enabled air handling unit resulted in frequent contamination of tissue culture samples. A handheld air sampler (SAS Super 100) was used, which could sample 100 liters of air per minute. Tryptic Soy Agar and Sabouraud dextrose agar plates were used to sample bacteria and fungi, respectively from the air. A fixed volume of air was sampled using the bio-sampler. Plates were placed in and removed from the bio-sampler in an aseptic manner. Plates were incubated at 25±2 ^0^C (for fungal cultivation) and 37±2 ^0^C (for bacterial cultivation) for 48 hours. Post-incubation, the number of colonies appeared were enumerated and converted to CFU/m^3^ using statistical conversion provided by the manufacturer. Control plates were used to ensure the sterility of the entire process.

## Acknowledgments

The authors acknowledge the Electron Microscopy Facility at Bangalore Life Science Cluster (C-CAMP, NCBS, inStem) for technical assistance in Scanning Electron microscopy imaging. Financial assistance was received from Department of Biotechnology-Biotechnology Industry Research Assistance Council (DBT-BIRAC), Govt. of India under Small Business Innovation Research Initiative (SBIRI) (Grant no. BT/SBIRI1372/31/16), COVID-19 Consortium (Grant no. BT/COVID0025/01/20), and from Karnataka Innovation and Technology Society (KITS) Department of Electronics, IT, Bt and S&T, Government of Karnataka, under Ideas2PoC Grant.

## Author contributions

KSP, DK carried out experiments designed and supervised by SD, AG. DM, NH carried out theoretical and numerical work supervised by JA, GT. JV, KSR, SD, AG conceptualized and designed the technology. JV, AG secured and managed funding.

## Competing interests

JA, GT declare no competing interests. SD is Director on Biomoneta board. The rest of the authors are, or were, employees of Biomoneta Research Private Limited, Bangalore, India 560065.

## Supplementary information

### S1. Range of microbes

To analyze microbe’s motion in laminar flow between electrode-plates we adopt the following approximations: (1) The flow is identical to that between infinitely wide plates, which is justified because *W/H*≫ 1, where *H* is the gap between electrode-plates and *W* is their width (perpendicular to the flow direction); (2) The flow is fully developed, which is justified because *L/H*≫ 1, where *L* is the length of electrode-plates along the flow direction; (3) The microbes move with the flow except when electric force acts on them, which is justified because the Stokes number of microbes (a measure of its inertial response to changes in the flow) is ≪ 1; (4) Self weight of microbes is negligible compared to the electric forces acting on them, because of their extremely small size (*<* 5 *µ*m).

The orientation of our coordinate system is shown in figure 1. A steady, unidirectional, incompressible, fully-developed flow is governed by [1]:

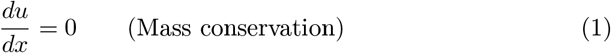

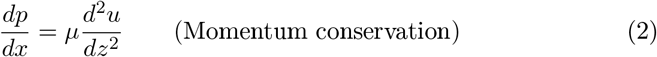

where *u* is the flow velocity along *X* direction, *p* is pressure and *µ* is dynamic viscosity of the fluid. Since, subject to our assumptions, *u* depends only on *z*, mass conservation in Eqn. 1 is automatically satisfied. Since the flow is induced by imposing a pressure difference between the ends of the electrode-plates, the pressure gradient *dp/dx* is a constant. Therefore, momentum conservation in Eqn. 2 is satisfied if *u* is a quadratic function of *z*. We assume it to be of the form, *u* = *Az*(*H − z*), because this automatically satisfies the no-slip boundary condition on the electrode-plates located at *z* = 0, *H*. The constant *A* is determined by computing the volumetric flow rate and equating it to the known value 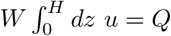, which yields:

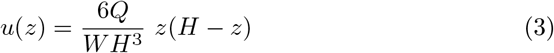

where *Q* is the volumetric flow rate of air between the electrode-plates. The flow velocity varies only along *Z* direction, being maximum midway between the plates and zero at the plates themselves (no-slip condition). Because the Reynolds number of microbe’s motion is ≪, 1, due to its small size and small speeds, only Stokes drag force is exerted by the ambient fluid, *F*_drag_ = 6*πµwa*, where *µ* is the dynamic viscosity of air. We have assumed that the microbe can be approximated by an equivalent sphere of radius *a*. The drag counterbalances the electric force on the microbe, *F*_electric_ = *qE*, where *q* is the surface charge on the microbe and *E* is the strength of the applied electric field. Equating the two forces yields for the settling velocity:

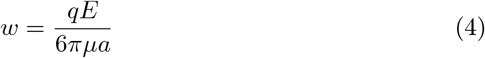

While drifting towards the electrode-plate the microbe also travels a distance *R* in the flow direction, which we call its “range”, refer figure 1. If *z*_0_is the initial distance of the microbe from the attracting electrode-plate at its entrance *x* = 0, then the time *T* needed for the microbe to hit the electrode-plate is, *T* = *z*_0_*/w*. After time *t*, the vertical location of the microbe initially located at *z*_0_ will be *z* = *z*_0_ − *wt*. Then, from eqn. 3, the microbe’s streamwise speed at that time will be *u*(*z*_0_ − *wt*). The microbe will hit the electrode-plate in time *T* = *z*_0_*/w* (assuming sufficient plate length). Therefore the range of the microbe beginning at location *z*_0_ is given by 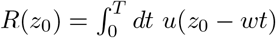 Changing the integration variable to *z* = *z*_0_ − *wt* transforms the integral to: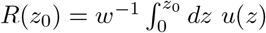. Substituting from eqn. 3 and integrating yields:

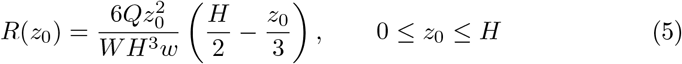

The microbe that is farthest from the attracting electrode-plate, i.e. at *z*_0_ = *H*, has the maximum range:

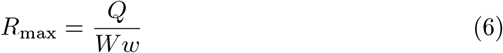

All the microbes entering ZeBox will hit the electrode-plate if its length is not less than the maximum range of the microbes, i.e. if *L*≥*R*_max_. Eq 5 for the range is visualized more easily if we divide it through by *R*_max_, Eq 6, and rewrite it in the following dimensionless form:

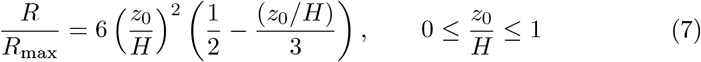

Assuming that all the trapped microbes are killed, Eq 7 plotted in supplementary figure 7 completely determines the microbicidal efficiency of ZeBox, as per the present model. Here, “microbicidal efficiency” is defined as the fraction of microbes entering electrode-plates that hit it and are inactivated, assuming a uniform distribution at the entrance. Using supplementary figure 7, the microbicidal efficiency is found as follows. We first compute *L/R*_max_ given the operating parameters. If *L/R*_max_≥ 1, then the microbicidal efficiency is of course 100%. Otherwise, we locate its value on the vertical axis of supplementary figure 7 and using the curve find the corresponding value on the horizontal axis, which gives the microbicidal efficiency. Therefore, *L/R*_max_ alone determines the microbicidal efficiency of ZeBox according to the present model.

**Supplementary figure 7:**
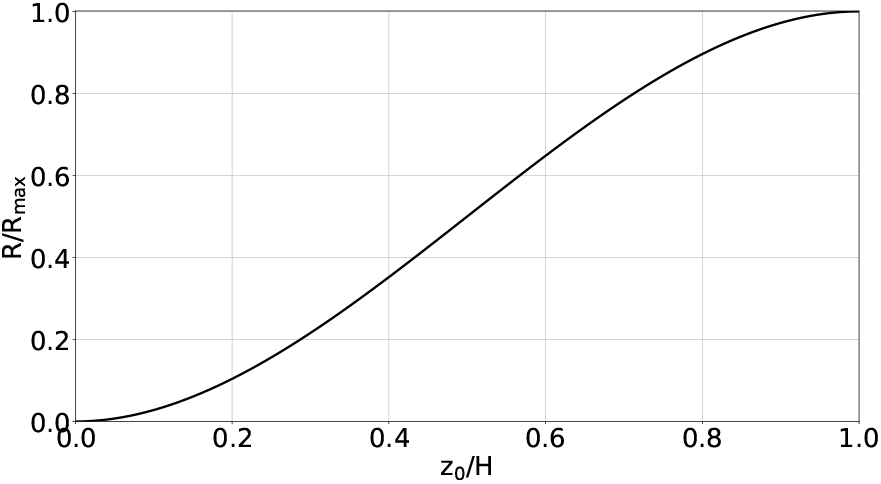
Range of a microbe. A microbe initially at distance *z*_0_*/H* from the attracting electrode-plate hits it at a distance *R/R*_max_.

### S2. Microbicidal efficiency of ZeBox

A microbe in an ionic solution is surrounded by a diffuse double layer of ions of molecular dimensions. The Debye length (*κ*^-1^), which is a measure of the thickness of the double layer, lies in the range: 1 *< κ*^*-*1^ *<* 10 nm [2]. Since the microbe’s size *a* ∼ 1 *µ*m, *κa*≫ 1 for microbes. The magnitude of the measured zeta potential (*ζ*) of microbes in phospate-buffer solution lies in the range 1 to 30 mV. Considering the worst-case-scenario, we may take *ζ* = 1 mV and *κa* = 100. The number of elementary charges *n* on the microbe may then be estimated as [2]:

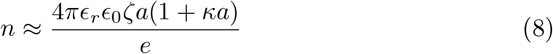

where *ϵ*_*r*_ is the dielectric constant of the solution, *E*_0_ = 8.9 × 10^*-*12^ F/m is the permittivity of vaccuum and *e* = 1.6 × 10^-19^C is the magnitude of the elementary charge. Eqn. 8 is derived from the linearized Poisson-Boltzmann equation governing the variation of electric potential due to distribution of ions in the diffuse layer surrounding a charged sphere; the linearization is a consequence of the Debye-Hückel approximation which is applicable when zeta potential is small (compared to 25 ∼ mV at 25 ^0^C) [2]. For a measured dielectric constant of 78.5 for the buffer solution, Eqn. 8 reveals *n >* 5000 elementary charges. Even allowing for an order-of-magnitude error and assuming *n >* 500 instead, Eqn. 4 yields a settling velocity of *w >* 7 cm/s, for *E* = 3 kV/cm in air.

Since the flow rate between a pair of electrode-plates is *Q <* 3 cfm and the electrode-plate width *W* = 10.9 cm, Eqn. 6 shows that *R*_max_ *<* 19 cm. In comparison, ZeBox employs 30 cm long electrode-plates. Although in theory it implies 100 % microbicidal efficiency for ZeBox, the present model is only approximate because it does not account for the effects of possible turbulence in the flow and slippage of microbes on the surface. In reality, as mentioned in the Results section, we obtain 83-99 % microbicidal efficiency for ZeBox as deduced from the measured microbial load reduction.

